# The impact of the genetic background on gene deletion phenotypes in *Saccharomyces cerevisiae*

**DOI:** 10.1101/487439

**Authors:** Marco Galardini, Bede P. Busby, Cristina Vieitez, Alistair S. Dunham, Athanasios Typas, Pedro Beltrao

## Abstract

Loss-of-function (LoF) mutations associated with disease don’t manifest equally in different individuals, a phenomenon known as incomplete penetrance. The impact of the genetic background on incomplete penetrance remains poorly characterized. Here, we systematically assessed the changes in gene deletion phenotypes for 3,786 gene knockouts in four *Saccharomyces cerevisiae* strains and 38 conditions. We observed 16% to 42% of deletion phenotypes changing between pairs of strains with a small fraction conserved in all strains. Conditions causing higher WT growth differences and the deletion of pleiotropic genes showed above average changes in phenotypes. We further illustrate how these changes affect the interpretation of the impact of genetic variants across 925 yeast isolates. These results show the high degree of genetic background dependencies for LoF phenotypes.

## Introduction

While a mutation can be associated with specific disorders it has long been observed that not all individuals carrying the disease variant will manifest it. Even for diseases caused by mutations in a single gene (i.e. monogenic disorders) incomplete penetrance is frequent, presumably due to differences in the genetic background (Kammenga 2017; Hou et al. 2018). Modulators of penetrance of disease causing variants have been identified for many human diseases (Cohen et al. 2005; Flannick et al. 2014; Chen et al. 2016) as well as loss-of-function (LoF) mutations in different model organisms (Hamilton & Yu 2012; Chari & Dworkin 2013; Vu et al. 2015; Chow et al. 2016; Mullis et al. 2018). This impact of the genetic background on the phenotypic consequence of LoF mutations affects our ability to predict phenotypes based on genetic variants. In *S. cerevisiae* and *E. coli*, gene deletion phenotypes have been extensively measured for all genes across hundreds of stress conditions (Hillenmeyer et al. 2008; Nichols et al. 2011). However, genes carrying putative LoF mutations in different strains are only weakly predictive of expected gene deletion phenotypes (Jelier et al. 2011; Galardini et al. 2017; Wagih et al. 2018). Understanding the extent and the mechanisms by which the effect of LoF variants depend on the genetic background is critical for the development of personalized medicine.

While there are many known examples of background dependencies on LoF mutations few comprehensive studies have addressed this phenomenon. Studies in *S. cerevisiae* have shown that 5% of essential genes are dispensable between two closely related strains (Ryan et al. 2012) and that deletion phenotypes of 7 chromatin-associated genes show quantitative differences across 2 genetic backgrounds that could be mapped via segregant analysis (Mullis et al. 2018). In a systematic RNAi studies in *C. elegans*, 20% of the 1,400 genes tested had different mutant phenotypes across two backgrounds and natural variation in gene expression accounted for some of the observed differences (Vu et al. 2015). Recently, gene deletion libraries were generated for 3 other backgrounds of *S. cerevisiae* (Busby et al. 2018) other than the original reference lab strain library (Winzeler et al. 1999). Growth measurements of these knockout libraries in the presence of statin identified significant differences in gene deletion phenotypes across the 4 genetic backgrounds. The availability of these libraries allows for the systematic study of the impact of the genetic background on gene deletion phenotypes.

Here we have measured the condition specific growth phenotypes for 4 *S. cerevisiae* deletion collections in a panel of 38 perturbations. The phenotypes show a large variation across the genetic backgrounds with an average between 16 to 42% gene-condition phenotype associations changing in each strain across all pairwise comparisons. Genes with the largest number of strain dependent changes of phenotypes had above average number of genetic and physical interactions suggestive of a role of genetic interactions in these changes. Conditions eliciting variable growth rates among the wild-type strains tended to have also the largest number of strain specific variation in gene deletion phenotypes. Finally, we have measured the growth profiles of a panel of 1,006 *S. cerevisiae* natural isolates (Peter et al. 2018) across the same conditions, identifying several variants associated with differences in growth that we linked to causal genes through the gene deletion analysis.

## Results

### Condition specific gene deletion phenotypes for 38 conditions in 4 *S. cerevisiae* genetic backgrounds

We measured growth phenotypes for 17,186 total gene knockouts in 4 *S. cerevisiae* genetic backgrounds (S288C, UWOPS87-2421, Y55, YPS606) with 3786 gene deletions measured in all backgrounds. The four strains used are genetically diverse with an average of 5.4 to 5.9 SNPs/kb relative to the lab “reference” strain S288C (Winzeler et al. 1999; Busby et al. 2018, Figure 1A). The deletions were arrayed as colonies in a 1536 agar plate format and were robotically pinned onto agar plates containing the 38 different conditions (Methods, Figure 1B). Colony size at the endpoint was used as a proxy for fitness and deviation from the expected growth was calculated, taking account the replicate measurements, using the s-score (Collins et al. 2006; Kapitzky et al. 2010; Nichols et al. 2011, Figure 1B). Positive and negative s-scores indicate gene deletions that confer resistance and sensitivity to a given condition, respectively. The list and description of the 38 conditions is available in **Supplementary Table 1**. These include environmental stresses (e.g. heat, high osmolarity, DNA damage), drugs (e.g. caspofungin, clozapine), metabolic conditions (e.g. amino-acid starvation) or combinations of stressors.

**Figure 1.**
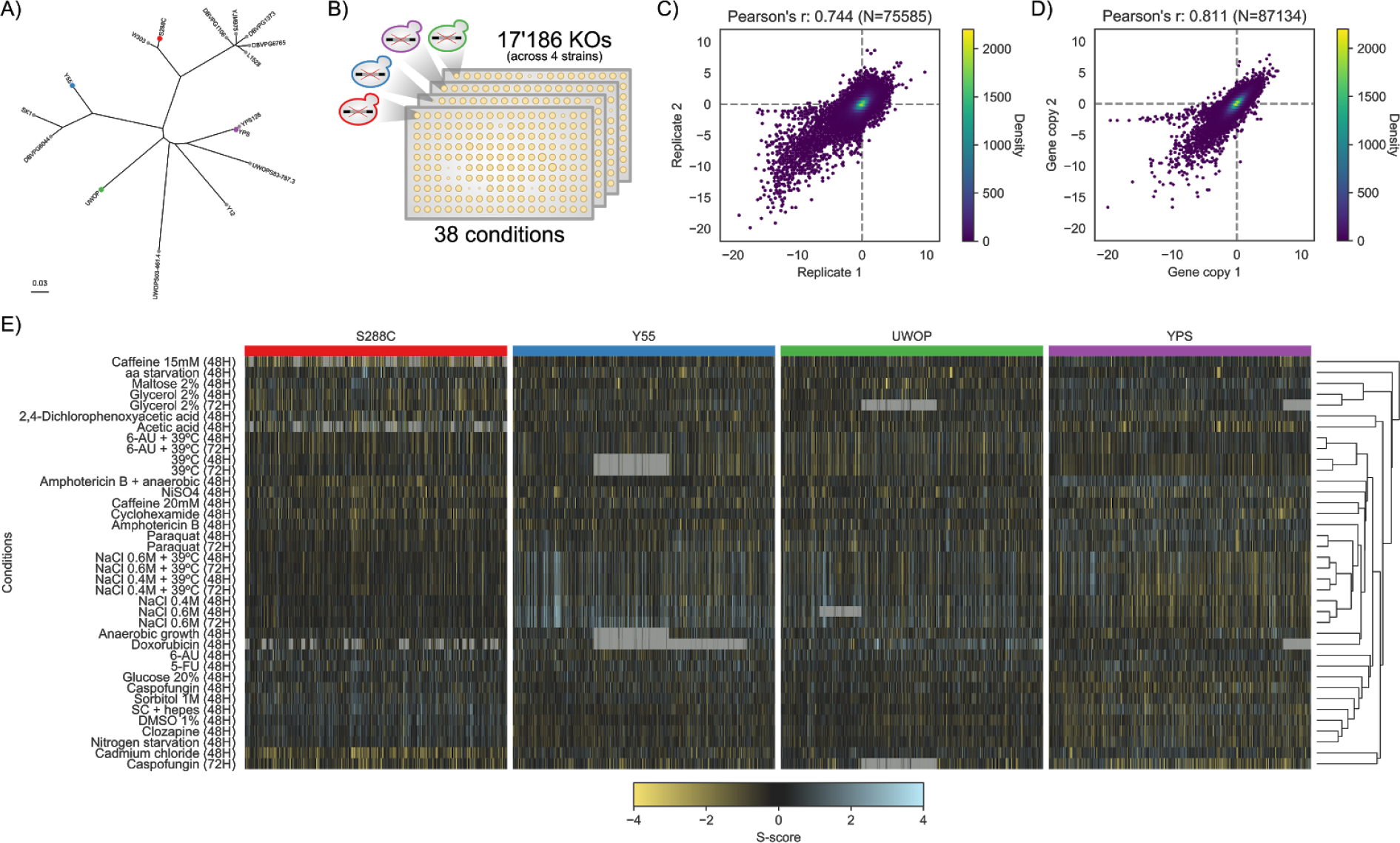
Chemical genomics screen across four S. cerevisiae strains. A) Core genome phylogeny of part of the Saccharomyces Genome Resequencing Project (SGRP) yeast isolates; coloured dots indicate the four strains whose KOs library was screened in this study. B) Schematic of the chemical genomics screen; each strain’s KO library was robotically plated on 1536 solid agar plates and each KO colony size was used as a proxy for fitness in each condition. C) Reproducibility of the S-scores using the two batches used in the screening. D) Reproducibility of the S-scores using genes having multiple independent colonies plated in the screening. E) Clustered heatmap of the whole chemical genomics screen; each subsection belongs to an individual strain’s KO library. Grey cells indicate missing values.

In total we measured 876,956 gene-condition phenotypic interactions that are provided as **Supplementary Material 1**. The assay used is highly reproducible as measured by the correlation of the s-scores using either 12 conditions that were replicated in 2 batches (Figure 1C, Pearson’s r=0.744 p-value<1E-50) or 2,293 genes that are spotted as replicates on the plates at different locations (Figure 1D, Pearson’s r=0.811 p-value<1E-50). The correlation of phenotypic scores for pairs of genes recapitulates known functional relationships (Supplementary Figure 1), further confirming the high quality of the screening data. We observed large differences in the profile of gene deletion phenotypes for the 4 different strains (Figure 1E) that can be quantified taking into account the high reproducibility of the assay.

### Quantification of genetic background differences of deletion phenotypes

The gene deletion phenotype scores for the 38 conditions were correlated across pairs of strains as a measure of similarity of their phenotypic profiles and plotted as a distribution for all genes in Figure 2A. On average the similarity of phenotype scores of the same gene knockout in different strains is only marginally higher than the observed for the correlation of deletion phenotype scores for random pair of genes (Figure 2A). The lack of correlation could be explained by the large fraction of KOs with no strong response across the four strains, resulting in differences in quantitative scores dominated by technical variability. In line with this, the similarity of phenotypes across strains increases for gene knockouts having larger numbers of significant deletion phenotypes (Figure 2A).

**Figure 2.**
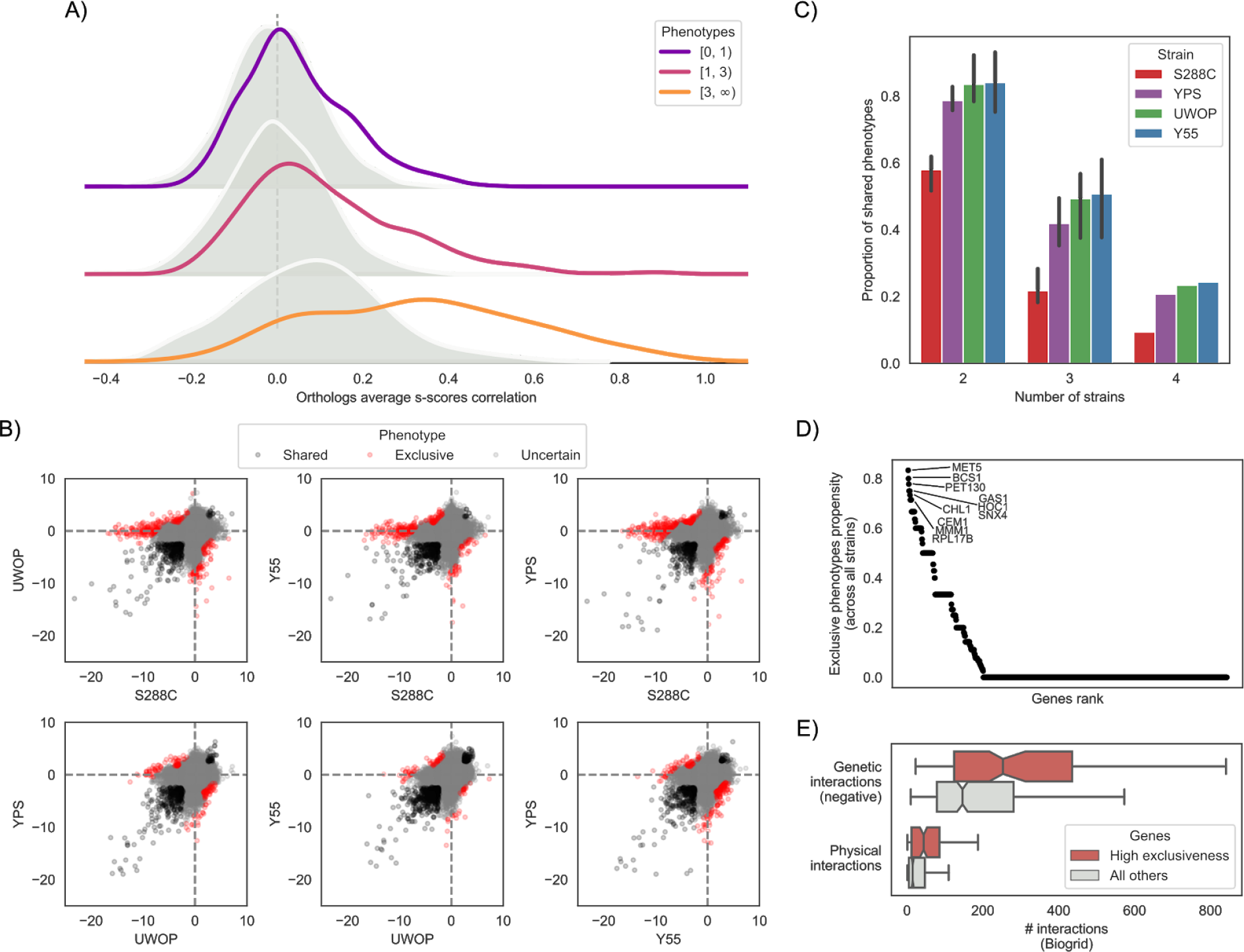
Systematic assessment of genetic background dependencies of gene deletion phenotypes. A) Average S-score Pearson’s correlation between the same genes (orthologs, solid line) and random gene pairs (shaded distribution) across all the 38 conditions and four strains. Genes are stratified by the number of conditions in which they show a significant phenotype across the four strains. B) S-score scatterplots for each pairwise strain comparison, highlighting conserved phenotypes (black points), significant changes (red points) and gene-condition relationships for which no call can reliably be made (grey points). C) Fraction of deletion phenotypes in each strain conserved with other stains in pairwise, three-way and four-way comparisons. D) Gene exclusiveness: a measure of each gene propensity to change its chemical genomics profile across strains. The top 10 genes’ names are reported. E) Genes with high exclusiveness (> 0) tend to have a higher number of negative genetic and physical interactions (as reported in the biogrid database).

In order to identify statistically significant differences of phenotypes we used an empirical null model that takes into account the variance of the assay and the mean dependence of the variance for the s-score (Bandyopadhyay et al. 2010, Methods, Supplementary Figure 2). For each pair of strains we identify the gene deletion phenotypes that were significantly shared (Figure 2B, black) or exclusive (Figure 2B, red) to each genetic background. Even though a part of the observed changes might be false positives, we are confident that the homogeneity in experimental conditions as well as excluding uncertain cases from the analysis (**Methods**) helps reducing these cases to a small number. We then used these significantly shared/exclusive phenotypes, excluding all other phenotypes (Figure 2B, grey), to calculate the fraction of shared/exclusive phenotypes for each pair of strains. We performed all pairwise comparisons and for each strain we then calculated the average fraction of shared phenotypes with the other 3 strains (Figure 2C), which ranged from 58% for S288C to 84% for Y55. This fraction drops further for phenotypes significantly conserved across more strains with 22% to 51% observed in 3 strains and 9% to 24% of gene-deletion phenotypes significantly conserved in all 4 backgrounds (Figure 2C). These highly conserved phenotypes include very central genes relevant for the corresponding responses such as sensitivity to osmotic stress (ΔHOG1), drug efflux (ΔPDR5) and amino acid biosynthesis (ΔADE, ΔMET, ΔSER, ΔTRP), among others (**Supplementary Table 2**). A small fraction (3% to 5%) of the phenotype that are not conserved between pairs of strains show a reversal in sign whereby the deletion causes resistance to the stress in one background but increased sensitivity in another background (**Supplementary Table 3**). We observed the strongest reversal for ΔMET5 exposed to amino acid starvation, which has a strong sick phenotype in YPS and UWOP, but shows increased resistance when knocked out in S288C. Since the S288C KO library is based on the BY4741 ΔMET17 (Cherry et al. 2012) strain, a potential explanation for this phenotype reversal could be a positive genetic interaction between MET5 and MET17. We observed few changes in these proportions when varying the significance threshold for calling phenotypes, indicating the robustness of this trends (Supplementary Figure 3).

We next focused on the gene knockouts that had the largest number of background dependent changes in phenotypes. We ranked all gene deletions according to the proportion of changes over all tested phenotypes (Figure 2D, **Supplementary Table 4**) and observed that the genes that change their deletion phenotypes at least once (N=242) had also a higher number of genetic and physical interactions partners based on data collected in the Biogrid database (Chatr-Aryamontri et al. 2017) for the S288C lab strain (Figure 2E). This suggests that gene pleiotropy is correlated with the probability that a deletion phenotype will depend on the genetic background. These genes were also enriched in GO terms related to endosomal transport (q-value=0.006), mitochondrion organization (q-value=0.007), cellular respiration (q-value=0.007) and cell wall organization or biogenesis (q-value=0.009, **Supplementary Table 5**). These genes were not more likely than others to show expression changes across the 4 yeast strains (Fisher’s exact test p-value > 0.05, **Supplementary Material 2**).

### Condition specific wild-type growth differences contribute to variance in gene deletion phenotypes

For each condition, we counted the number of changes in deletion phenotypes observed across genetic backgrounds (Figure 3A and Supplementary Figure 4). We compared these changes with the average growth rate differences of the 4 WT strains (i.e. no knockout) in the same conditions (Figure 3B and Supplementary Figure 4). We noticed that some conditions having large number of changes corresponded to cases where a significant growth difference was observed for the WT strains. For example, S288C grows poorly under maltose relative to the other strains (Chow et al. 1989) (Figure 3B) and also showed the largest number of exclusive gene deletion phenotypes (Figure 3A), the same was observed for caffeine. In contrast, the high salt (osmotic shock) conditions, S288C had the highest WT growth and the smallest number of phenotype differences. While this was not the case for all conditions there is a significant trend where a slower WT strain growth in a condition is associated with a larger number of strain specific knockout phenotypes (Figure 3C, Pearson’s r=-0.21, p-value=0.02).

**Figure 3.**
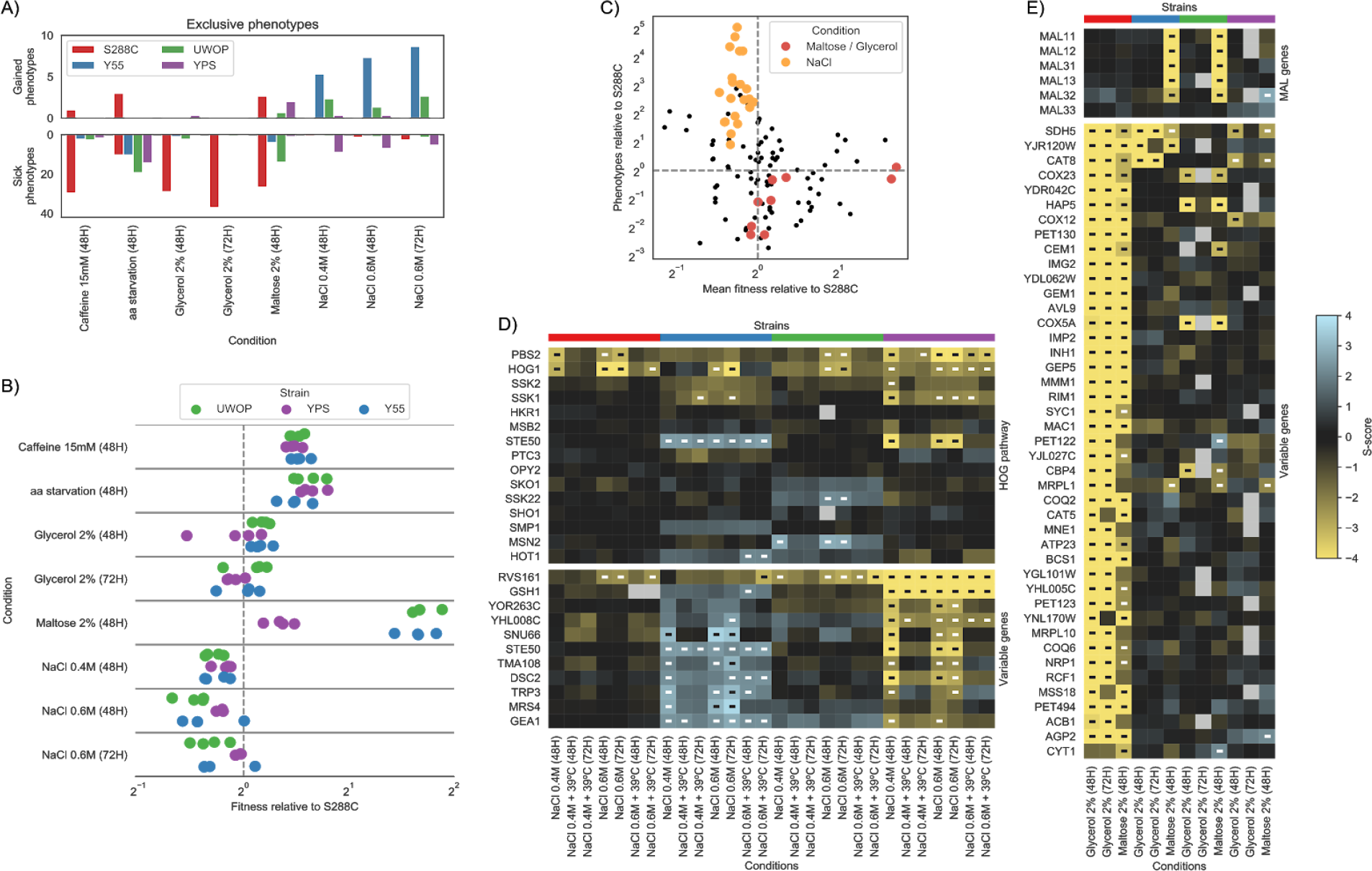
Wild type condition specific growth differences are associated with the degree of gene deletion phenotype differences. A) Barplots reporting the average number of gain and sick phenotypes that are specific to each strain across all pairwise comparisons. B) Wild type fitness of each strain relative to S288C across the same conditions as in panel A; each dot represents a specific replicate where colony sizes were measured. C) Relationship between the wild type fitness relative to S288C and the number of conditionally essential genes relative to S288C; each dot represents a strain-condition replicate as in panel B. D) Changes in gene deletion phenotypes for growth on osmotic stress conditions. The top heatmap contains genes belonging to the HOG pathway, while the bottom one those genes whose growth phenotypes varies the most between Y55 and YPS. E) Conditional gene essentiality changes for growth on glycerol and maltose. The top heatmap contains the MAL genes, while the bottom one those genes whose growth phenotypes vary the most between S288C and the other three strains.

We analyzed in more detail some conditions with large changes in phenotypes. For high salt, that elicits an osmotic stress, deleting the two central kinases of the osmotic shock pathway (HOG1 and PBS2) generally impaired growth in all backgrounds, as expected. This pathway has two upstream branches converging on PBS2/HOG1 (Brewster et al. 1993; Hohmann 2009). These branches can be redundant and thus show few phenotypes under osmotic stress. However, the STE50 deletion shows striking differences causing increased sensitivity in YPS background, resistance in the Y55 and no phenotype in S288C and UWOP. Similarly to STE50, we identified 10 more genes with strain specific phenotypes in high salt condition (Figure 3D). Some of these genes are related to osmosensing (STE50, RVS161), ER function (DSC2, GEA1) and metabolism (MRS4, TRP3).

For growth under maltose the two strains with the best WT growth (UWOP and Y55) also had strong growth defects when maltose induced genes, present in two clusters, were deleted (Figure 3E). It is known that S288C does not grow effectively in maltose due to inactivation of the maltose activator proteins in the MAL loci (Chow et al. 1989). It is therefore expected that deleting MAL genes (i.e. maltose induced genes) causes no further decrease in growth in maltose negative strains such as S288C but has a strong impact on MAL positive strains such as Y55 and UWOP. Similar to S288C, the YPS strain appears to also be a maltose negative strain. The poor growth under maltose then creates additional vulnerabilities to the cell, rendering essential a large number of genes involved in non-fermentable growth under maltose for S288C (Figure 3E). Interestingly the same set of genes are required for S288C to grow in glycerol, suggesting that S288C should grow poorly under glycerol, although this did not translate in a strong growth defect of the WT under this condition (Figure 3B).

### Quantitative trait analysis of condition specific growth differences in a panel of 1,006 *S. cerevisiae* strains

To test whether our the gene deletion phenotypes data in different genetic backgrounds could be used to better understand the impact of natural variation on phenotypes, we performed a quantitative trait analysis (QTL) across 47 conditions using a panel of 925 *S. cerevisiae* natural isolates (i.e. the fraction of the tested 1,006 strains with available genotype data, Peter et al. 2018). Growth measurements, fitness measurements and phenotype calculations were performed as for the deletion libraries (**Methods**). The s-score measurements used (Figure 4A and **Supplementary Figure 5**) represents, as above, condition specific growth phenotype for each strain where a genetic background can specifically affect growth under a given condition. Using the genomic variants we identified a total of 151,673 common single nucleotide polymorphisms (SNPs, minor allele frequency > 5%). In addition, we predicted the impact of missense variants in each coding region and calculated a probability of LoF for each gene in each strain (Jelier et al. 2011; Galardini et al. 2017); this could be regarded as a gene disruption or gene burden score. We then performed a QTL analysis for each condition using the common SNPs, the gene burden score, the gene copy-number variation (CNV) and the presence/absence patterns of genes as predictors (**Methods**). In total we found 579 significant associations, with the largest number of associations observed for growth under Amphotericin B and caffeine (Figure 4B), both known to have an impact on the cell wall. Both conditions are also unlikely to be present in the natural environment and therefore genetic variants causing growth defects under these conditions are less likely to be selected against. Common SNPs had the highest number of significant associations (365, 0.005%), followed by gene presence/absence (159, 0.805%), gene burden score (29, 0.047%) and CNVs (26, 0.060%).

**Figure 4.**
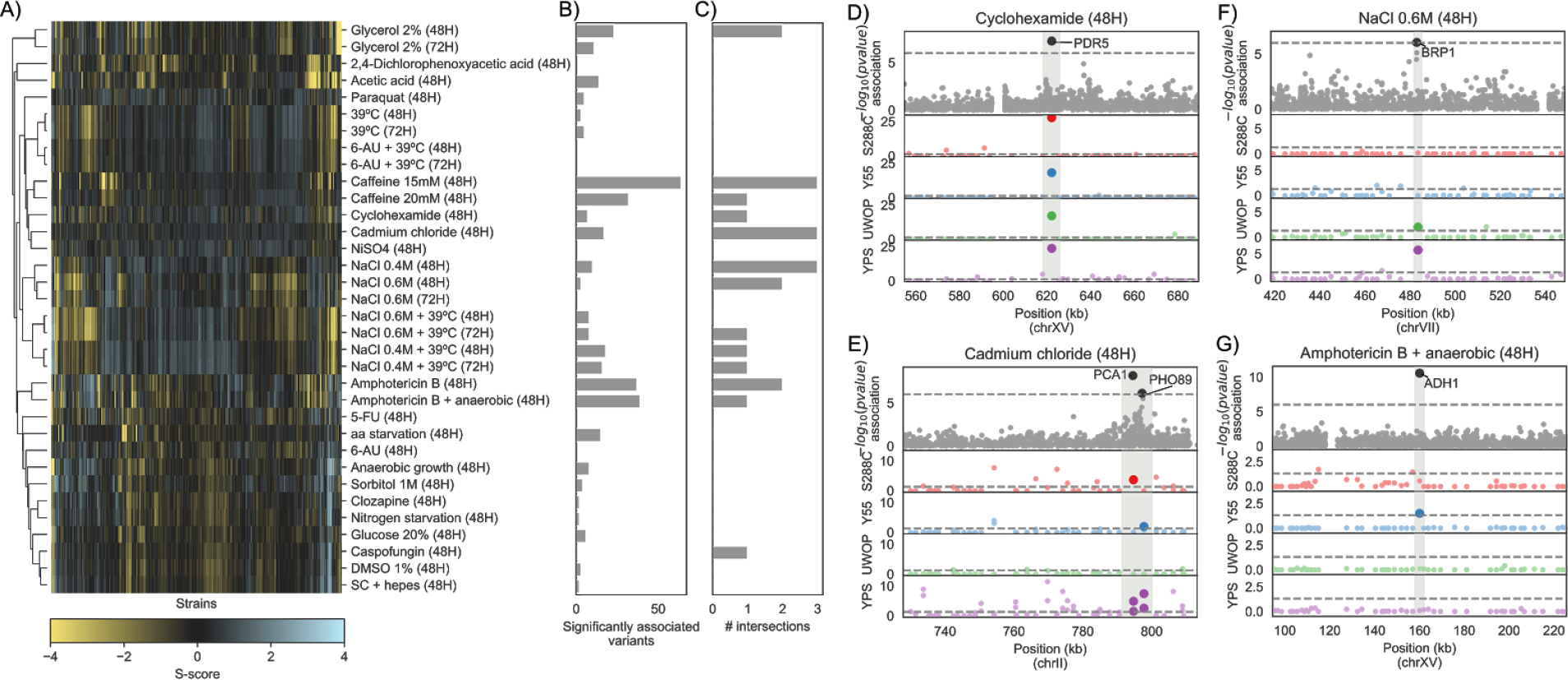
Genes linked to growth phenotypes via QTL analysis in 925 natural yeast isolates. A) S-score heatmap of the yeast natural isolates across 34 conditions that were also used in the KO screening. B) Number of variants significantly associated (p-value < 1E-6) with phenotypic variation in each growth condition. C) Number of associated variants that overlap (i.e. are in a 3kbp window) with a conditionally essential gene in the same condition, in any of the four KO libraries. D-G) Manhattan plots showing examples of overlaps between associated variants and the KO screening. The top plot shows the associations between variants and growth in the natural isolates as a function of the −log_10_ of the association p-value, while the four bottom plots show the strength of the KO phenotypes across the four yeast strains, as a function of the −log_10_ of the corrected s-score p-value. Sections shaded in grey indicate the overlap between associations and KO data. Position in the yeast chromosome is reported in kilobase units.

For each condition we obtained a list of genes associated with growth differences from the gene deletion analysis and crossed it with the variants, and their linked genes, associated with growth differences across the 925 yeast strains. Unexpectedly, we found no significant enrichment between the gene-condition associations obtained from the QTL analysis and the gene-condition associations found in the gene deletion experiments (Fisher’s exact test p-value > 0.05). Despite the lack of overall enrichment, several QTL associations can be validated by the gene deletion information (Figure 4C, **Supplementary Table 6**). For example, 141 strains had a high gene burden score in the PDR5 locus, which had a significant association with growth in the presence of Cyclohexamide. Deletion of this ABC transporter is known to cause multidrug resistance and showed Cyclohexamide dependent deletion phenotypes in all 4 backgrounds (Figure 4D). The presence of two SNPs in two other transporters, cadmium transporting P-type ATPase (PCA1) and a membrane Na+/Pi cotransporter (PHO89) were linked to growth under cadmium chloride and had also significant gene deletion phenotypes in at least 3 of the strains (Figure 4E). A SNP close to BRP1 showed an association with growth under high salt stress which is supported by BRP1 deletion phenotypes in 2 strains. However, several other cases had fewer support. For example, the absence of the ADH1 gene in 450 strains showed an association with anaerobic growth in the presence of amphotericin B. Deletion of this gene shows a strong phenotype for this condition only when knocked out in Y55. We found a total of 22 gene~condition associations overlapping between the QTL and KO analysis with most overlaps observed with gene deletion phenotypes exclusive to a single genetic background.

## Discussion

Our results show that the genetic background has a strong impact on gene deletion phenotypes in *S. cerevisiae*. The fraction of significant differences across two individuals (18 to 40%) is similar to the fraction of changes observed for RNAi phenotypes for two strains of *C. elegans* (20%, Vu et al. 2015). The fraction of shared phenotypes decreases further for the common set observed across all strains (N=73) with typically <25% significantly shared phenotypes across the 4 strains. Analysis of additional backgrounds would be needed to fully access the fraction of gene deletion phenotypes that is independent of the genetic background.

The large-scale analysis allowed us to search for general trends associated with the observed differences. Strains having a slower WT growth in a given condition also tended to have a larger number of gene deletion phenotypes in those conditions, suggesting that in such conditions the poor growing strains have more modes of failure and are impacted by a larger number of gene deletions. Growth in maltose serves as a good example of how existing genetic variation can interact with LoF mutations. The S288C strain, has genetic variants that cause it to not be able to grow well in maltose and therefore, in this condition, becomes reliant on genes required for non-fermentable growth. It remains a challenge to be able to find similar justifications for how the genetic background interacts with the gene deletions for other conditions but those identified here could be further studied using a segregant analysis as previously done by Mullis and colleagues (Mullis et al. 2018).

Some genes had a higher proportion of changes of their deletion phenotypes. These tended also to have an above average number genetic and physical interactions. The interaction assays used as the basis for this analysis have been conducted in the S288C background strain but they nevertheless likely reflect the degree of pleiotropy of each gene. One interpretation of this result would be that genes that are involved in multiple processes are more likely to have also larger number of changes in deletion phenotypes since there will be many ways by which the genetic background difference may interact with the loss-of-function of these genes. Of the four strains, the reference strain (S288C) stands out as the one where specific gene-condition associations are the most abundant when compared to the other three strains: 58% versus ~80%. This observation combined with other idiosyncrasies specific to this strain (Mortimer & Johnston 1986; Winston et al. 1995; Brachmann et al. 1998), such as growth in the presence of maltose, indicates that observations made on a domesticated individual might not necessarily reflect natural populations.

Lastly, we performed a QTL analysis using 925 strains for the same conditions and using the same experimental set-up. We were expecting that variants associated with differential growth in a given condition were linked to genes that also caused gene deletion phenotypes in the same condition. Overall we found no such enrichment which may relate to the fact that natural isolates are likely to have few LoF variants, in particular associated to conditions that are experienced in the environment. The total number of strains used is likely also to be very limiting since there are typically very few strains showing strong growth differences across any specific condition. Larger number of strains or segregant analysis (Bloom et al. 2013; Cubillos et al. 2013) could be used in the future to further study the relationship between natural variants and deletion phenotypes. Despite an overall lack of enrichment, our results suggest that interpretation of the impact of genetic variants using the gene deletion information available for a single genetic background is unlikely to be comprehensive.

In summary our results suggest that interpretation of the impact of genetic variants on the phenotypes of individuals would likely need detailed gene-phenotype information in more genetic backgrounds than that of a model individual.

## Methods

### Strains used

Mata haploid KO libraries in the genetic backgrounds of S288C (Winzeler et al. 1999, 4’889 KOs), UWOPS87-2421 (abbreviated as UWOP, 4’014 KOs), Y55 (4’190 KOs) and YPS606 (Busby et al. 2018, abbreviated as YPS, 4’093 KOs) were used to assess if different genetic backgrounds have an effect on gene deletion phenotypes. These libraries were maintained on YPD+G418 prior to screening in 384 colony format. The 1’006 natural isolate strain collection (a kind gift from Gianni Liti, Peter et al. 2018), was maintained on YPD in 384 colony format prior to screening.

### Chemical genomic analysis

Growth of KO libraries, and 925 strain collection was evaluated on concentrations of chemical and environmental stress conditions (**Supplementary Table 1**) that inhibit the growth of S288C by approximately 40%. The libraries were maintained and pinned with a Singer RoTor in 1536 colony format. Synthetic complete (Kaiser et al. 1994) media was used with or without the stress condition, incubated at 30°C (unless temperature was a stress) for 48h or 72h, and imaged using a SPIMAGER (S&P Robotics) equipped with a Canon Rebel T3i digital camera.

The growth measurements were performed in three separate batches with overlapping sets of conditions to judge for variation of the method. The first batch of the chemical genomic screening was carried out with the S288C deletion collection and used the following conditions: anaerobic, amphoteracin B, nystatin, DMSO, 2,4,D, glycerol, maltose, hepes buffered medium, caffeine, 6-AU, paraquat, 39°C, sorbitol. Batch two was carried out with the deletion collections in the four genetic backgrounds (S288C, Y55, YPS606 and UWOPS87-2421) and contained all (13) of the conditions from batch 1 plus: 5-FU, doxorubicin, cadmium chloride, caspofungin, clozapine, Nickel sulfate, clioquinol, high glucose (20%), minimal medium, nitrogen starvation medium, cyclohexamide, sodium chloride 0.4M and 0.6 M, and the duel stress conditions sodium chloride 0.4 and 0.6M plus 39°C, 6-AU plus 39°C and amphoterician B plus anaerobic growth. The two batches were carried out as separate experiments.

### Chemical genomics data analysis

Raw plate images were cropped using ImageMagick to exclude the plate plastic borders. Raw colony sizes were extracted from the cropped images using gitter (Wagih & Parts 2014) v1.1.1, using the “autorotate” and “noise removal” features on. Poor quality plates were flagged when no colony size could be reported for more than 5% of colonies (poor overall quality) or when no colony size could be reported for more than 90% of a whole row or column (potential grid misalignment); known empty spots in each plate where used to flag incorrect plates. Overall less than 5% of all pictures have been discarded (175/4221, 4.15%). Conditions with less than three replicates across the two experiment batches were excluded from further processing. Raw colony sizes for the remaining conditions were used as an input for the EMAP algorithm (Collins et al. 2006), with default parameters except the minimum colony size which was set to 5 pixels. The algorithm computes an S-score, which indicates whether the growth of each KO is deviating from the expected growth in each condition. The raw s-scores were further quantile normalized in each condition. Significant loss-of-function and gain-of-function phenotypes were highlighted by transforming the s-scores in z-scores, given that the s-scores in each condition follow a normal distribution. P-values were derived using the survival function of the normal distribution and corrected using an FDR of 5% (false discovery rate). The whole dataset, comprising 876,956 gene-condition interactions is available as **Supplementary Material 1**.

The overall relative fitness of the three non-reference strains (Y55, UWOP and YPS) against the S288C reference was computed as follows:

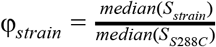

where *median* (*S*) is the median normalized colony size. The normalized colony size is computed in each plate by first applying a surface correction step, followed by a border correction step. The surface correction is applied to reduce the impact of spatial abnormalities on colony sizes; in short, the second-degree polynomials of the row and column indices are computed in a matrix, which is then qr factorized. The resulting matrix is used to construct an ordinary least squares linear model between the *Q* factorized matrix and the corresponding vector of raw colony sizes. The surface normalized colony sizes are then computed as follows:

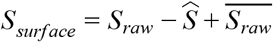

where 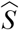 is the size prediction from the ordinary least squares model. The surface corrected colony sizes (*S*_*surface*_) are further corrected to take into account the border effect, meaning the difference in size between colonies in the two outermost rows and columns with respect of the rest of the plate. The border correction is computed as follows:

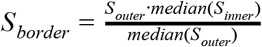

where *median* (*S*_*outer*_) is the median size of colonies on the outer border of the plate and *median* (*S*_*inner*_) is the median size of colonies in the rest of the plate. The overall fitness is computed only in those cases where the plates belonging to the four strains’ KO libraries have been screened at the same time, in order to make the comparison robust to changes in experimental conditions. The relative fitness measures are available in **Supplementary Table 7**.

To test whether the four KO libraries are able to recapitulate known gene functional relationships, we tested if gene pairs belonging to the same functional groups tended to have correlated S-score vectors in each of the four yeast strains. Two functional relationships sets were used: the CYC2008 protein complexes set (Pu et al. 2009) and Kegg modules belonging to *S. cerevisiae* (Muto et al. 2013). The KOs common to all four libraries were selected and for each strain only those that had at least one phenotype with corrected p-value below 0.01 were used to compute the Pearson correlation of S-scores between each gene pair. The ability of these gene-gene correlations to recapitulate the known functional relationships was assessed by constructing Receiver Operator Characteristic and Precision-Recall curves, using the known relationships as the true positives set and 10 random gene pairs sets with same numerosity as the true set as true negative set.

The conservation or similarity of gene deletion phenotypes across the four yeast strains was assessed computing the Pearson’s correlation between the s-score profiles across all conditions of the same genes in all pairs of strains and then computing the average of these values. Genes were stratified by the average number of conditions in which they show either a loss-of-function or a gain-of-function phenotype across the four strains. Random gene pairs were used as background.

### Reproducibility of chemical genomics screens

The reproducibility of the chemical genomics screen was assessed in two separate ways. The first method assessed the technical reproducibility of the s-scores across the two batches in which the screen was conducted. The raw pictures were divided according to the batch of origin and the EMAP algorithm (Collins et al. 2006) was used to compute a set of S-scores for each batch. For the 13 conditions that were tested in both batches the S-score correlation was computed. We refer to this analysis as both technical and biological because the inoculates are derived from the same source plate but at very different times (**Supplementary Material 3**). Biological replicability was assessed using 2,293 KOs that are pinned exactly twice across the library.

### Significant changes in conditional gene essentiality

Significant changes in chemical genomics profiles between any two strains were computed following a previously published approach that also used s-scores (Bandyopadhyay et al. 2010). The two set of s-scores computed as part of the batch replicate analysis were used as a null model for the absence of changes in s-scores, as a way to estimate the degree of expected variation observed across different experiments. Since the variance in S-scores is higher at higher absolute s-score values, this has to be taken into account when calling significant differences; a sliding window approach was applied when constructing the null model. Given the two set of s-scores the following vectors were computed:

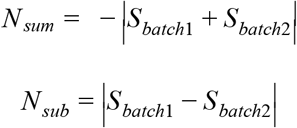

where *S*_*batch*1_ and *S*_*batch*2_ are the S-scores from the replicate batches, respectively. The sliding window was then applied to *N*_*sum*_, dividing the vector in 100 slices with at least 20 observation in each one and recording the mean 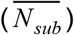 and standard deviation (σ_*sub*_) of *N*_*sub*_ for each slice. For each strain pairwise comparison *N*_*sum*_ and *N*_*sub*_ were recorded for each matching slice and the corresponding 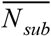 and σ_*sub*_ were extracted from the null distribution using a linear interpolation. A normal distribution with mean 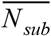 and standard deviation σ_*sub*_ was then constructed around each slice and the cumulative distribution of the normal function was used to derive a p-value to indicate significant differences. The p-value was FDR corrected and incoherent differences were assigned a corrected p-value of 1; specifically, those cases where both strains have a significant phenotype but corrected p-value lower < 0.01 (150 comparisons over 875,833) and cases where both strains do not show a significant phenotype but corrected p-value lower < 0.01 (26 comparisons over 875,833). The full dataset comprising 875,834 comparisons is available as **Supplementary Material 4**.

When looking at the proportion of significant loss-of-function or gain-of-function phenotypes that each strain shares with the other strains, we considered those comparisons were the focal strain had a significant gain-of-function or loss-of-function phenotype and corrected p-value < 0.01 (phenotype not shared) and were both strains had a significant phenotype and corrected p-value >= 0.01 (shared phenotype); all other comparisons were not considered.

Each gene’s propensity to change its conditional essentiality profile across the six strains’ pairwise comparisons (*E*, exclusive phenotype propensity) was computed as follows:

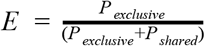

where *P*_*exctusive*_ is the number of loss-of-function or gain-of-function phenotypes that vary significantly (corrected p-value < 0.01), while *P*_*shared*_ is the number of loss-of-function or gain-of-function phenotypes in both strains of the comparison that do not vary significantly (corrected p-value >= 0.01). A gene whose *E* > 0 was considered with “high exclusiveness” (**Supplementary Table 4**).

Variable genes in Figure 3D and 3E were selected based on a corrected p-value cutoff of 1E-4 in at least one comparison.

### GO terms enrichment analysis

Gene gene ontology (GO) annotations were downloaded from the SGD database(Cherry et al. 2012), while the GO slim yeast dataset was downloaded from the gene ontology website (Ashburner et al. 2000; The Gene Ontology Consortium 2017). GO terms enrichments were assessed using goatools (Klopfenstein et al. 2018) v0.8.2, using a FDR-corrected p-value threshold of 0.01.

### Transcriptomics analysis

Yeast were grown to and OD of 0.4 and total RNA was extracted using the MasterPure Yeast RNA purification Kit (Biozym, Epicentre). The samples were quality tested on the Fragment analyzer (AATi/Agilent) using the Standard sensitivity RNA kit (AATi/Agilent), 600 ng of total RNA was used for library preparation. The libraries were prepared using the Truseq Stranded mRNA kit (Illumina) using a Beckman Fxp liquid handler system. Sequencing was carried out on an Illumina NextSeq 500 in 75 single end mode.

Raw single-ended Illumina reads were trimmed to remove the TruSeq3 adaptors using trimmomatic (Bolger et al. 2014) v0.36. Trimmed reads were pseudo-aligned to the yeast reference genome transcripts (downloaded from the SGD database, Cherry et al. 2012) using kallisto (Bray et al. 2016) v0.44.0 with the sequence based bias correction and using an average fragment length of 130bp (70bp standard deviation). Differential gene expression analysis between each each strain and S288C was performed using DESeq2 (Love et al. 2014) v3.8. Raw reads are available in the GEO database with accession number GSE123118.

### Natural isolates growth assay

Raw plate images were cropped using ImageMagick to exclude the plate plastic borders. Raw colony sizes were extracted from the cropped images using gitter (Wagih & Parts 2014) v1.1.1, using the “autorotate” and “noise removal” features on. Poor quality plates were flagged when no colony size could be reported for more than 5% of colonies (poor overall quality) or when no colony size could be reported for more than 90% of a whole row or column (potential grid misalignment). Conditions with less than three replicates were excluded from further processing. Raw colony sizes for the remaining conditions were used as an input for the EMAP algorithm (Collins et al. 2006), with default parameters except the minimum colony size which was set to 5 pixels. The computed raw s-scores were further analysed as reported in the “Chemical genomics data analysis” section.

### Genome-wide association study of the yeast natural isolates

The genetic variants found in the yeast natural isolates collection (SNPs/InDels, CNVs and genes presence/absence patterns) were downloaded from http://1002genomes.u-strasbg.fr/files/ on October 4th 2018. SNPs and InDels were normalized and filtered to retain variants with at least 5% minor allele frequency (common variants), using bcftools (Li et al. 2009) v1.9. Rare variants (SNPs with minor allele frequency <= 5%) were included computing their impact on gene function, using the “gene disruption score” described in previous studies (Jelier et al. 2011; Galardini et al. 2017). In short, common nonsynonymous and nonsense variants were kept, together with genes presence/absence patterns; the impact of nonsynonymous variants was predicted using the SIFT (Ng & Henikoff 2001) and FoldX (Guerois et al. 2002) algorithms, when applicable. The individual predictions were translated to their probability of being neutral (*P*_*neutral*_) based on a collection of variants with known impact (Jelier et al. 2011), using the following transformations:

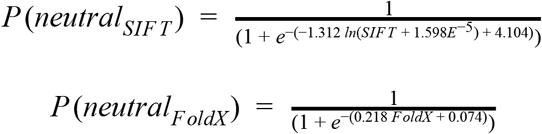

where *SIFT* and *FoldX* are the scores of each individual predictor. Nonsense variants were assigned a *P*_*neutral*_ value of 0.99 if they appeared in the last 5% of the protein sequence, 0.01 otherwise. The overall likelihood that gene function was affected by common variants (*P*(*AF*), or gene disruption score) was computed as follows:

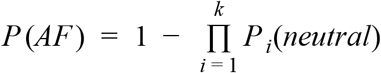

where *k* is the totality of nonsynonymous and nonsense variants observed in each gene. When both SIFT and FoldX predictions were available, priority was given to SIFT scores; nonsynonymous variants with neither predictions available were assigned the highest observed *P*_*neutral*_ value. If a gene was considered as absent a (*P*(*AF*) value of 0.99 was assigned. The gene disruption score is available in **Supplementary Material 5**. The CNVs, gene presence/absence patterns and digitized gene disruption scores (1 if (*P*(*AF*) > 0.9, 0 otherwise) were encoded in a VCF (Variant Calling Format) file together with the common variants and recoded using plink (Purcell et al. 2007) v1.90b4.

A genome-wide association analysis has been carried out to highlight common and rare variants associated with growth variability across the yeast natural isolates, using the LMM (Linear Mixed Model) implemented in limix (Lippert et al. 2014) v2.0.0a3. Missing values were mean imputed and the model was applied to each growth condition independently. The kinship matrix was computed using the strains’ phylogenetic tree from the original yeast natural isolates publication (Peter et al. 2018). Intersections between associated variants and genes present in the KO libraries were recorded using a 3kbp window centered around each gene. Gene annotations were retrieved from the SGD database (Cherry et al. 2012). Enrichments were tested using the Fisher’s exact test.

### Reproducibility

Most of the code used to process the data is available at the following URL: https://github.com/mgalardini/2018koyeast. The code is mostly based on the python programming language, using the following libraries: numpy (Oliphant 2006) v1.15.2, scipy (Oliphant 2007) v1.1.0, pandas (McKinney & Others 2010) v0.23.4, scikit-learn (Pedregosa et al. 2011) v0.20.0, statsmodels (Seabold & Perktold 2010) v0.9.0, biopython (Cock et al. 2009) v1.71, dendropy (Sukumaran & Holder 2010) v4.4.0. Reproducibility was ensured through the use of snakemake (Köster & Rahmann 2018) v4.7.0. Data was visualized inside jupyter notebooks using jupyter (Kluyver et al. 2016) v4.4.0 using the matplotlib (Hunter 2007) and seaborn (Waskom et al. 2018) plotting libraries version 3.0.0 and 0.9.0, respectively.

## Supporting information

## Acknowledgements

We thank Ferris Jung and Nayara Trevisan Doimo de Azevedo for the RNAseq work, Lars Steinmetz for the S288C deletion collection and Paul Atkinson for the Y55, YPS606 and UWOPS87-2421 deletion collections. Leopold Parts, Gianni Liti and Kevin Roy for critical reading of the manuscript.

## Supplementary figures

**Supplementary Figure 1.**
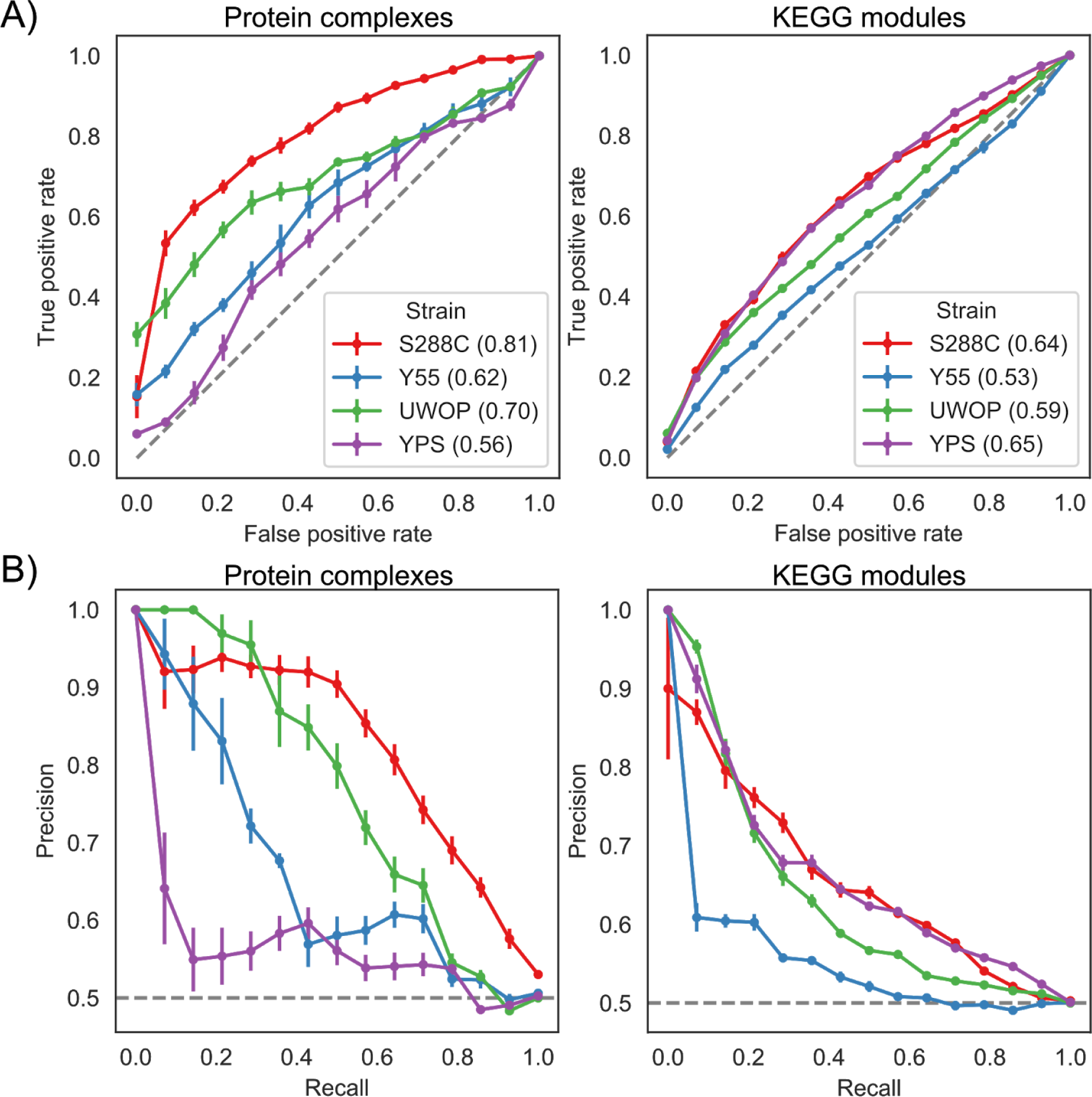
Benchmarking of s-scores. A) Receiver operating characteristic (ROC) curves and B) Precision-Recall curves for two gene functional interactions sets across the four KO libraries. Numbers between parentheses in A) represent the Area Under the Curve (AUC).

**Supplementary Figure 2.**
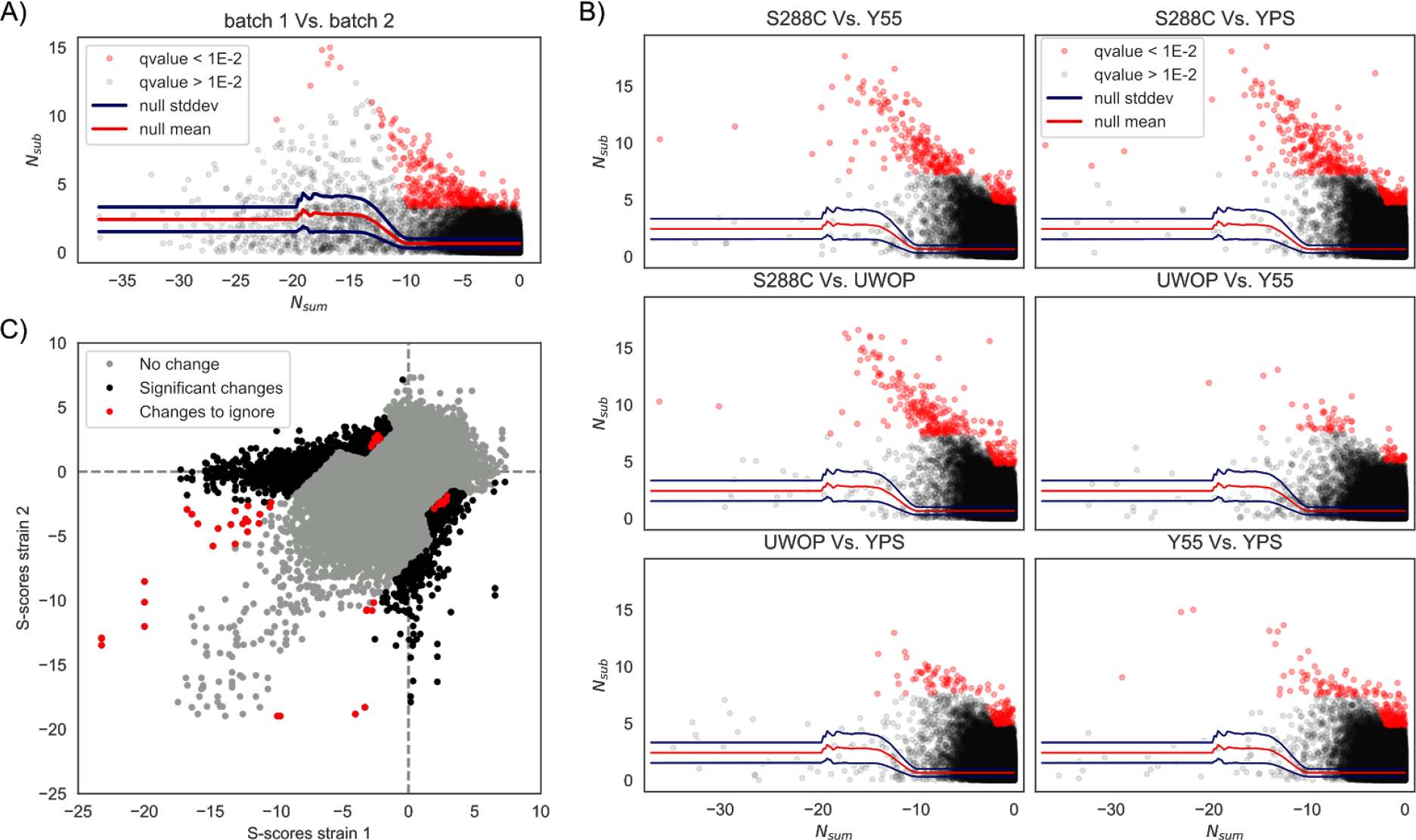
Calling significant differences in s-scores. A) Null model to call significant s-scores differences. B) All strains pairwise comparisons, indicating s-scores that change significantly between each pair. C) Overview of all strains pairwise comparisons; “changes to ignore” refers to significant changes (corrected p-value < 0.01) that are inconsistent (see Materials and methods).

**Supplementary Figure 3.**
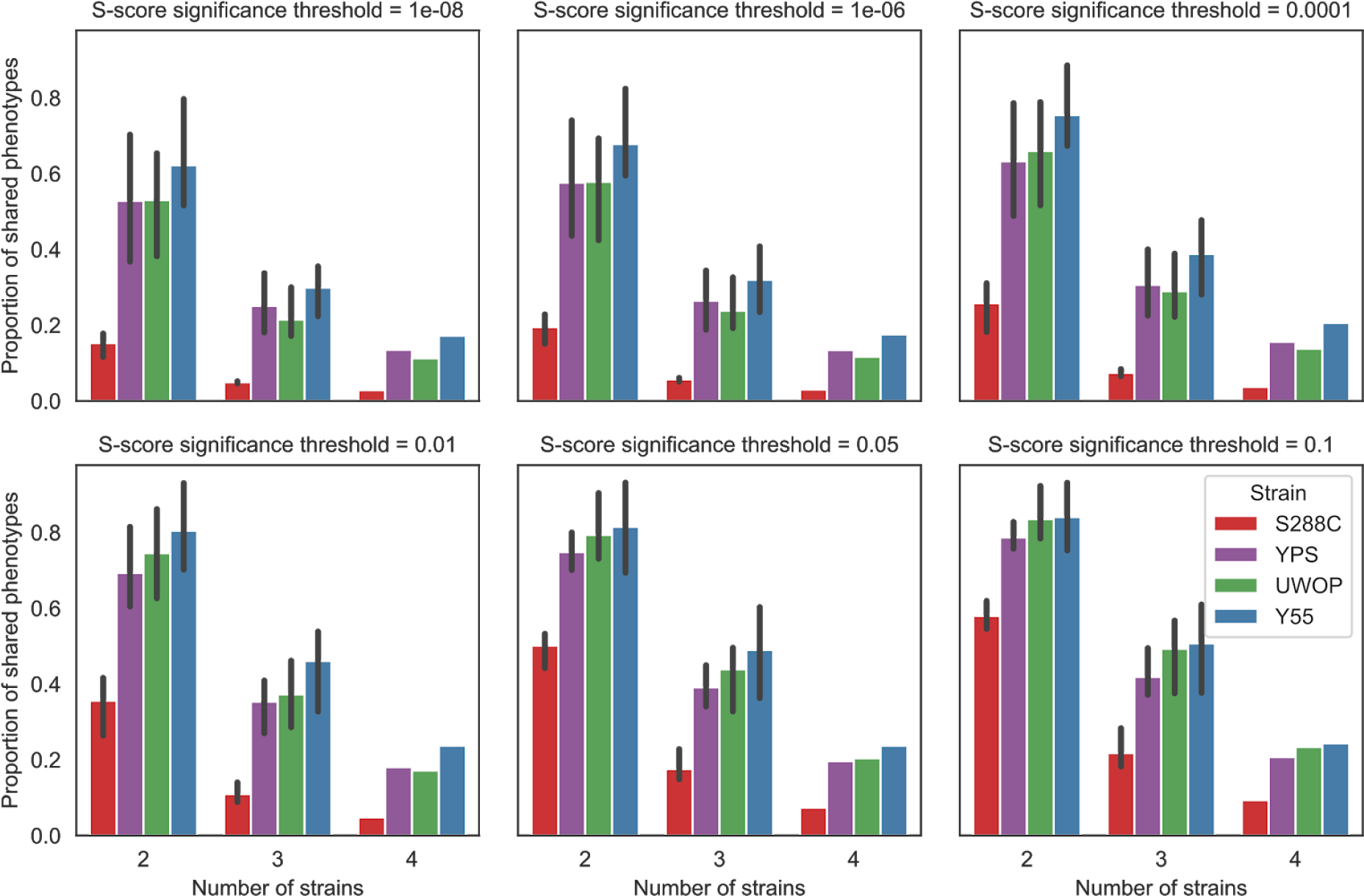
General invariance of shared phenotypes when varying the significance threshold. Proportion of shared phenotypes for different s-scores significance thresholds on calling phenotypes.

**Supplementary Figure 3.**
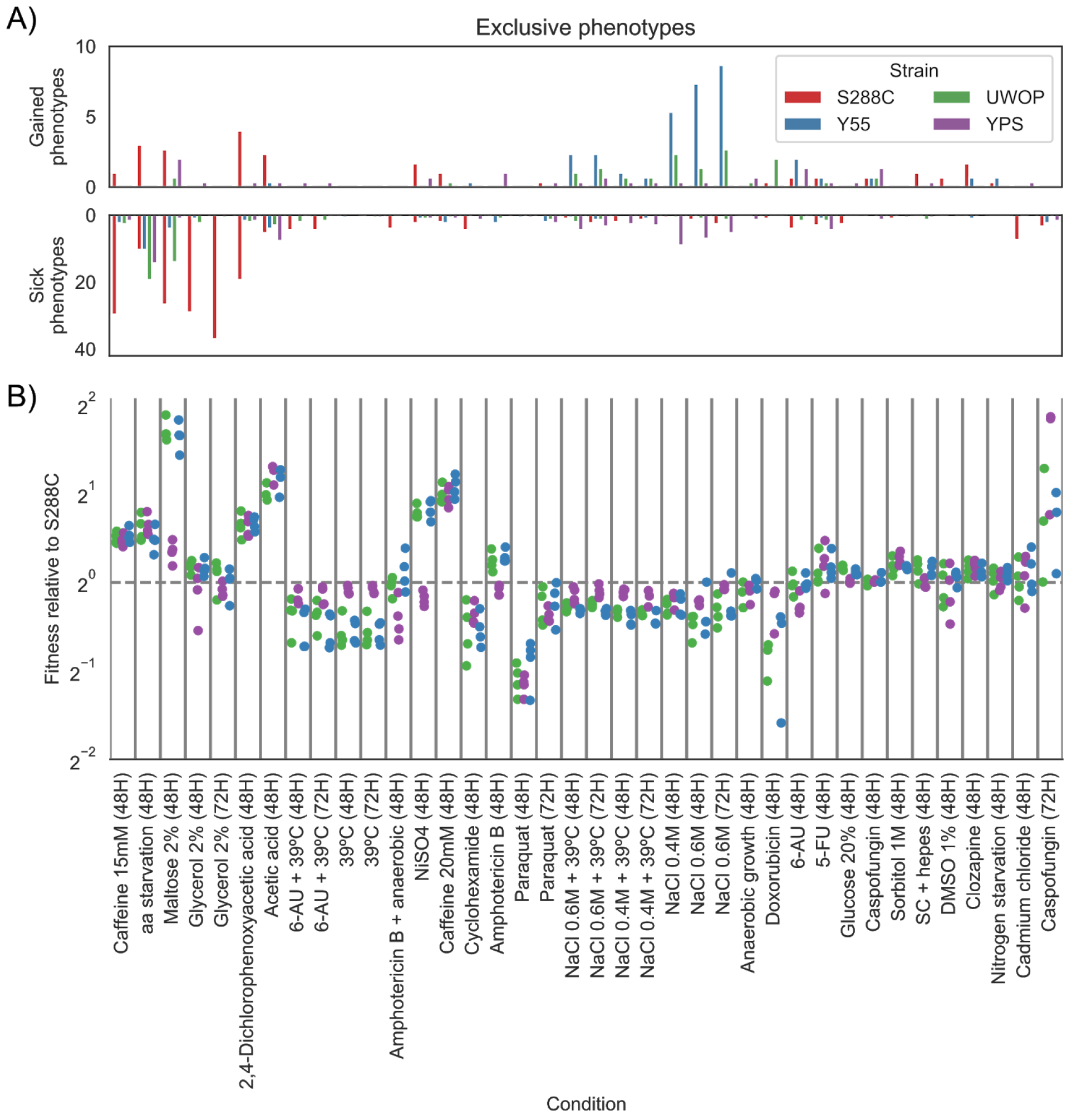
Condition specific changes in gene essentiality and fitness. A) Average number of KO phenotypes that are exclusive to a particular strain across all strains pairwise comparisons. Conditions order is the same as panel B. B) Wild type fitness of each strain relative to S288C across all conditions; each dot represents a specific replicate where colony sizes were measured.

**Supplementary Figure 4.**
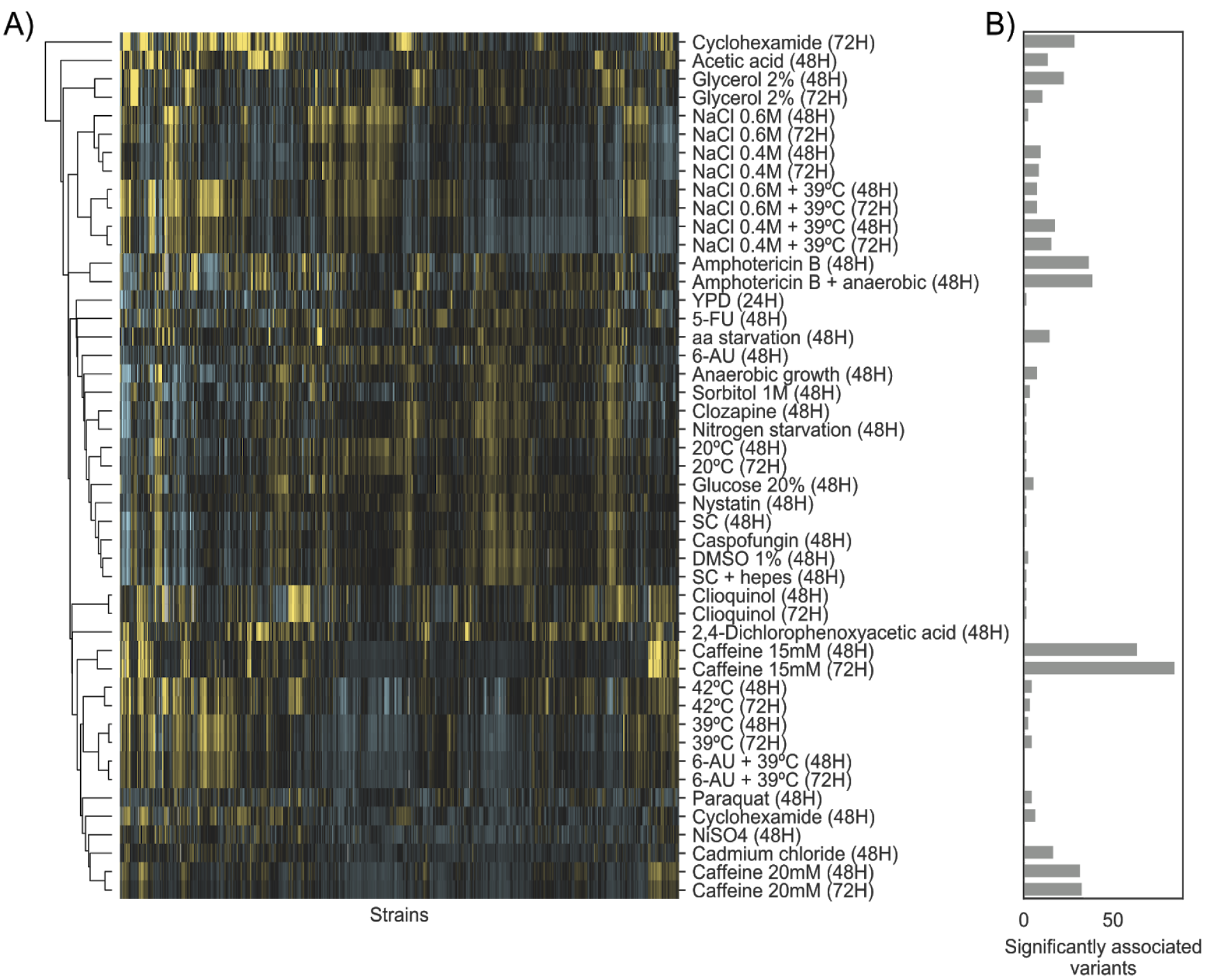
Growth profiles of the 925 yeast natural isolates. A) Clustermap of s-scores for the whole set of conditions in which the natural isolates were tested in. B) Number of significantly (p-value < 1E-6) associated variants across the same conditions as in panel A.

## Supplementary Materials

**Supplementary Table 1**: list of conditions used to profile the KO libraries

**Supplementary Table 2**: list of gene-condition associations conserved across the 4 KO libraries

**Supplementary Table 3**: list of gene-condition associations that show a phenotypes reversal between pairs of strains

**Supplementary Table 4**: exclusiveness measure for all genes tested for their ability to change their phenotype

**Supplementary Table 5**: GO terms enriched for genes with high exclusiveness

**Supplementary Table 6**: intersections between gene KO and QTL hits

**Supplementary Table 7**: fitness of the Y55, YPS and UWOP strains relative to S288C

**Supplementary Material 1**: KO libraries s-scores

**Supplementary Material 2**: differential expression of the Y55, YPS and UWOP strains compared to S288C in physiological conditions and in the presence of caffeine

**Supplementary Material 3**: KO libraries s-scores divided by batch in order to assess reproducibility

**Supplementary Material 4**: systematic assessment of changes in gene conditional essentiality for each possible strains’ pair

**Supplementary Material 5**: gene burden score for common variants in 1,011 strains across 5,699 genes

## Refrences

Ashburner, M. et al., 2000. Gene ontology: tool for the unification of biology. The Gene Ontology Consortium. Nature genetics, 25(1), pp.25–29.

Bandyopadhyay, S. et al., 2010. Rewiring of genetic networks in response to DNA damage. Science, 330(6009), pp.1385–1389.

Bloom, J.S. et al., 2013. Finding the sources of missing heritability in a yeast cross. Nature, 494(7436), pp.234–237.

Bolger, A.M., Lohse, M. & Usadel, B., 2014. Trimmomatic: a flexible trimmer for Illumina sequence data. Bioinformatics, 30(15), pp.2114–2120.

Brachmann, C.B. et al., 1998. Designer deletion strains derived from Saccharomyces cerevisiae S288C: a useful set of strains and plasmids for PCR-mediated gene disruption and other applications. Yeast, 14(2), pp.115–132.

Bray, N.L. et al., 2016. Near-optimal probabilistic RNA-seq quantification. Nature biotechnology, 34(5), pp.525–527.

Brewster, J.L. et al., 1993. An osmosensing signal transduction pathway in yeast. Science, 259(5102), pp.1760–1763.

Busby, B.P. et al., 2018. Master regulators of genetic interaction networks mediating statin drug response in Saccharomyces cerevisiae vary with genetic background. bioRxiv. Available at: https://www.biorxiv.org/content/early/2018/10/16/443879.abstract.

Chari, S. & Dworkin, I., 2013. The conditional nature of genetic interactions: the consequences of wild-type backgrounds on mutational interactions in a genome-wide modifier screen. PLoS genetics, 9(8), p.e1003661.

Chatr-Aryamontri, A. et al., 2017. The BioGRID interaction database: 2017 update. Nucleic acids research, 45(D1), pp.D369–D379.

Chen, R. et al., 2016. Analysis of 589,306 genomes identifies individuals resilient to severe Mendelian childhood diseases. Nature biotechnology, 34(5), pp.531–538.

Cherry, J.M. et al., 2012. Saccharomyces Genome Database: the genomics resource of budding yeast. Nucleic acids research, 40(Database issue), pp.D700–5.

Chow, C.Y. et al., 2016. Candidate genetic modifiers of retinitis pigmentosa identified by exploiting natural variation in Drosophila. Human molecular genetics, 25(4), pp.651–659.

Chow, T.H., Sollitti, P. & Marmur, J., 1989. Structure of the multigene family of MAL loci in Saccharomyces. Molecular & general genetics: MGG, 217(1), pp.60–69.

Cock, P.J.A. et al., 2009. Biopython: freely available Python tools for computational molecular biology and bioinformatics. Bioinformatics, 25(11), pp.1422–1423.

Cohen, J. et al., 2005. Low LDL cholesterol in individuals of African descent resulting from frequent nonsense mutations in PCSK9. Nature genetics, 37(2), pp.161–165.

Collins, S.R. et al., 2006. A strategy for extracting and analyzing large-scale quantitative epistatic interaction data. Genome biology, 7(7), p.R63.

Cubillos, F.A. et al., 2013. High-resolution mapping of complex traits with a four-parent advanced intercross yeast population. Genetics, 195(3), pp.1141–1155.

Flannick, J. et al., 2014. Loss-of-function mutations in SLC30A8 protect against type 2 diabetes. Nature genetics, 46(4), pp.357–363.

Galardini, M. et al., 2017. Phenotype inference in an Escherichia coli strain panel. eLife, 6, pp.1–19.

Guerois, R., Nielsen, J.E. & Serrano, L., 2002. Predicting changes in the stability of proteins and protein complexes: a study of more than 1000 mutations. Journal of molecular biology, 320(2), pp.369–387.

Hamilton, B.A. & Yu, B.D., 2012. Modifier genes and the plasticity of genetic networks in mice. PLoS genetics, 8(4), p.e1002644.

Hillenmeyer, M.E. et al., 2008. The chemical genomic portrait of yeast: uncovering a phenotype for all genes. Science, 320(5874), pp.362–365.

Hohmann, S., 2009. Control of high osmolarity signalling in the yeast Saccharomyces cerevisiae. FEBS letters, 583(24), pp.4025–4029.

Hou, J. et al., 2018. Genetic Network Complexity Shapes Background-Dependent Phenotypic Expression. Trends in genetics: TIG, 34(8), pp.578–586.

Hunter, J.D., 2007. Matplotlib: A 2D Graphics Environment. Computing in Science Engineering, 9(3), pp.90–95.

Jelier, R. et al., 2011. Predicting phenotypic variation in yeast from individual genome sequences. Nature genetics, 43(12), pp.1270–1274.

Kaiser, C., Michaelis, S. & Mitchell, A., 1994. Methods in yeast genetics. Available at: http://genesdev.cshlp.org/content/9/3/local/back-matter.pdf.

Kammenga, J.E., 2017. The background puzzle: how identical mutations in the same gene lead to different disease symptoms. The FEBS journal, 284(20), pp.3362–3373.

Kapitzky, L. et al., 2010. Cross-species chemogenomic profiling reveals evolutionarily conserved drug mode of action. Molecular systems biology, 6(451), p.451.

Klopfenstein, D.V. et al., 2018. GOATOOLS: A Python library for Gene Ontology analyses. Scientific reports, 8(1), p.10872.

Kluyver, T. et al., 2016. Jupyter Notebooks -- a publishing format for reproducible computational workflows. In F. Loizides & B. Schmidt, eds. Positioning and Power in Academic Publishing: Players, Agents and Agendas. IOS Press, pp. 87–90.

Köster, J. & Rahmann, S., 2018. Snakemake-a scalable bioinformatics workflow engine. Bioinformatics, 34(20), p.3600.

Li, H. et al., 2009. The Sequence Alignment/Map format and SAMtools. Bioinformatics, 25(16), pp.2078–2079.

Lippert, C. et al., 2014. LIMIX: genetic analysis of multiple traits. bioRxiv, p.003905. Available at: https://www.biorxiv.org/content/early/2014/05/22/003905 [Accessed November 22, 2018].

Love, M.I., Huber, W. & Anders, S., 2014. Moderated estimation of fold change and dispersion for RNA-seq data with DESeq2. Genome biology, 15(12), p.550.

McKinney, W. & Others, 2010. Data structures for statistical computing in Python. In Proceedings of the 9th Python in Science Conference. pp. 51–56.

Mortimer, R.K. & Johnston, J.R., 1986. Genealogy of principal strains of the yeast genetic stock center. Genetics, 113(1), pp.35–43.

Mullis, M.N. et al., 2018. The complex underpinnings of genetic background effects. Nature communications, 9(1), p.3548.

Muto, A. et al., 2013. Modular architecture of metabolic pathways revealed by conserved sequences of reactions. Journal of chemical information and modeling, 53(3), pp.613–622.

Ng, P.C. & Henikoff, S., 2001. Predicting deleterious amino acid substitutions. Genome research, 11(5), pp.863–874.

Nichols, R.J. et al., 2011. Phenotypic landscape of a bacterial cell. Cell, 144(1), pp.143–156.

Oliphant, T.E., 2006. A guide to NumPy, Trelgol Publishing USA.

Oliphant, T.E., 2007. SciPy: Open source scientific tools for Python. Computing in Science and Engineering, 9, pp.10–20.

Pedregosa, F. et al., 2011. Scikit-learn: Machine Learning in Python. Journal of machine learning research: JMLR, 12(Oct), pp.2825–2830.

Peter, J. et al., 2018. Genome evolution across 1,011 Saccharomyces cerevisiae isolates. Nature, 556(7701), pp.339–344.

Purcell, S. et al., 2007. PLINK: a tool set for whole-genome association and population-based linkage analyses. American journal of human genetics, 81(3), pp.559–575.

Pu, S. et al., 2009. Up-to-date catalogues of yeast protein complexes. Nucleic acids research, 37(3), pp.825–831.

Ryan, O. et al., 2012. Global gene deletion analysis exploring yeast filamentous growth. Science, 337(6100), pp.1353–1356.

Seabold, S. & Perktold, J., 2010. Statsmodels: Econometric and statistical modeling with python. In Proceedings of the 9th Python in Science Conference. SciPy society Austin, p. 61.

Sukumaran, J. & Holder, M.T., 2010. DendroPy: a Python library for phylogenetic computing. Bioinformatics, 26(12), pp.1569–1571.

The Gene Ontology Consortium, 2017. Expansion of the Gene Ontology knowledgebase and resources. Nucleic acids research, 45(D1), pp.D331–D338.

Vu, V. et al., 2015. Natural Variation in Gene Expression Modulates the Severity of Mutant Phenotypes. Cell, 162(2), pp.391–402.

Wagih, O. et al., 2018. Comprehensive variant effect predictions of single nucleotide variants in model organisms. bioRxiv.Available at: https://www.biorxiv.org/content/early/2018/05/02/313031.abstract.

Wagih, O. & Parts, L., 2014. gitter: a robust and accurate method for quantification of colony sizes from plate images. G3, 4(3), pp.547–552.

Waskom, M. et al., 2018. mwaskom/seaborn: v0.9.0 (July 2018), Available at: https://zenodo.org/record/1313201.

Winston, F., Dollard, C. & Ricupero-Hovasse, S.L., 1995. Construction of a set of convenient Saccharomyces cerevisiae strains that are isogenic to S288C. Yeast, 11(1), pp.53–55.

Winzeler, E.A. et al., 1999. Functional characterization of the S. cerevisiae genome by gene deletion and parallel analysis. Science, 285(5429), pp.901–906.

